# The genome of the avian malaria parasite *Haemoproteus majoris* (lineage WW2) and its relationship to other *Plasmodium* species

**DOI:** 10.64898/2026.07.13.738139

**Authors:** Amélie Bardil, Arnaud Berthomieu, Jacques Dainat, Michael C. Fontaine, Olof Hellgren, Ana Rivero, Thomas D. Otto, Sylvain Gandon

## Abstract

Avian malaria parasites form a highly prevalent and genetically diverse group within the haemosporidians, yet they have long been overlooked relative to their human- and rodent-infecting counterparts. Among these, parasites of the genus *Haemoproteus* (Haemosporida, Haemoproteidae) are widespread and prevalent blood parasites of birds, transmitted by louse flies (Hippoboscidae) and biting midges (Ceratopogonidae). Recent phylogenomic analyses place *Haemoproteus* parasites at the root of the haemosporidian tree, making genomic data from these taxa essential for understanding the evolutionary origins of malaria parasites. To date, only two avian *Plasmodium* and one avian *Haemoproteus* genomes have been sequenced. We present the first assembled genome of *Haemoproteus majoris* (lineage WW2), a common blood parasite of passerine birds. As avian erythrocytes are nucleated, parasite DNA was enriched by FACS-based sorting to discriminate and isolate the parasite from host cell nuclei prior to whole-genome amplification. The genome was assembled using Nanopore long-read sequencing and polished with Illumina short-reads, yielding 145 contigs with a total assembly size of 23.9Mb and a G+C content of 27.85%. Genome annotation identified 5501 protein-coding genes, 69 non-coding RNA genes, and 57 long terminal repeat retrotransposons (LRT-RTs), including one full-length element. This genomic resource represents a critical step towards elucidating the evolutionary history and genomic architecture of avian malaria parasites.

**SIGNIFICANCE STATEMENT:** We present the first assembled genome of *Haemoproteus majoris* (lineage WW2), a prevalent and generalist avian malaria parasite. Taxonomic resolution of this genus is difficult as there are few distinct morphological differences among closely-related species. This genome provides a valuable resource for studying the evolution within the *Haemoproteus* genus and to elucidate the evolutionary history of malaria parasites.

## INTRODUCTION

Although malaria parasites are best known for infecting humans, they also parasitize hundreds of other vertebrate species, including non-human primates, reptiles, and birds. Avian malaria, in particular, represents one of the most prevalent, widespread, and genetically diverse groups of haemosporidian parasites, comprising four genera: *Plasmodium*, *Leucocytozoon*, *Haemoproteus*, and *Parahaemoproteus*, each containing several hundred distinct cytochrome-b lineages [1]. While multiple high-quality genomes are available for human and rodent malaria *Plasmodium* parasites, genomic resources for other avian haemosporidians remain scarce. This limits our ability to conduct comparative genomics studies involving population genetics or gene evolution in this otherwise well studied host-parasite system. Sequencing avian malaria genomes is particularly challenging because avian red blood cells are nucleated, which complicates the isolation of parasite DNA free from host contamination. As a result, our ability to explore the genetic diversity of this highly diverse and ecologically significant group is hindering both comparative genomic analyses and deeper insights into their evolutionary biology. Despite this, two high-quality genomes of avian Plasmodium have been assembled using paired-end Illumina-sequencing, enabled by different parasite DNA enrichment approaches: *Plasmodium relictum* (lineage pSGS1-like/DONANA05) [1,2] and *Plasmodium gallinaceum* (strain 8A) [2]. These assemblies comprise 498 and 152 scaffolds, with total sizes of 22.6Mb and 23.8Mb, GC contents of 18.34% and 17.83%, and *N*_50_ values of 1,833,711bp and 1,343,538bp, respectively. In comparison, the only published *Haemoproteus* genome, that of *Haemoproteus tartakovskyi* (lineage SISKIN1), is a highly fragmented 454 pyrosequencing assembly of 2,983 scaffolds totaling 23.2Mb, with a GC content of 25.4%, a longest scaffold of 108,686bp and a *N*_50_ of 17,240bp [3].

Here we present the first high quality assembled genome of *Haemoproteus majoris* lineage WW2, a biting midge-transmitted blood parasite, and one of the most prevalent and generalist *Haemoproteous* species in European passerine birds. *H. majoris* has been found to infect more than 30 bird species across 10 different families, reaching prevalences of up to 50% in some populations [4]. Recent phylogenomic analyses place *Haemoproteus* parasites at the root of the haemosporidian tree, making genomic data from these taxa essential for understanding the evolutionary origins of malaria parasites [5]. This genome therefore constitutes a valuable addition to the scarce genomic resources available for avian haemosporidians, providing a foundation for comparative genomic analyses across the haemosporidian tree, and for placing the evolutionary origins of malaria parasites in a broader phylogenomic context.

## RESULTS AND DISCUSSION

### Sequencing and genome assembly

Whole genome sequencing yielded 390,964 Nanopore raw reads (depth: 158X) and 501,165,175 Illumina paired-end raw reads (depth: 6295X). As neither the plastid nor the mitochondrial genomes could be recovered, all analyses focus on the nuclear genome.

The nuclear genome of *H. majoris* WW2 is similar in size and GC content to that of the published *H. tartakovskyi* SISKIN1 (Table 1) [3]. However, our assembly represents a substantial improvement in contiguity as it comprises only 145 contigs and no ambiguous bases (N’s) compared to the 2,983 scaffolds reported for the SISKIN1 lineage [3] and the 498 scaffolds reported for *P. relictum* SGS1-like [2]. This improvement is attributable to the combined use of deep long- and short-read sequencing. A total of 5,570 genes were identified in *H. majoris* WW2 (Table 1), comparable to *H. tartakovskyi* SISKIN1 (5,988 genes), and to other *Plasmodium* species [2,3]. Of these, 5,501 were protein-coding, 11 were putative ribosomal RNA (rRNA) genes, and 40 were putative transfer RNA (tRNA) genes.

**Table 1:** Assembly and annotation statistics of the nuclear genome of *H. majoris* lineage WW2, compared to two of the previously published avian malaria genomes: *H. tartakovskyi*, lineage SISKIN1 and *P. relictum* lineage SGS1-like (renamed DONANA05, used for gene annotation and orthology analysis). Statistics for SISKIN1 genome the scaffold (HtScaffold0932) which contains the complete 5,992 bp mitochondrial genome [2]. Transposable elements have not been characterized for SISKIN1.

| **Nuclear genome assembly statistics** | ***H. majoris***<br>**WW2** | ***H. tartakovskyi***<br>**SISKIN1** | ***P. relictum***<br>**SGS1-like** |
| --- | --- | --- | --- |
| Genome size (Mb) | 23.9 | 23.2 | 22.6 |
| N50 | 363,479 | 17,240 | 1,833,711 |
| GC content (%) | 27.85 | 25.4 | 18.34 |
| Assembly level | contig | scaffold | pseudo-chromosome |
| No. of contigs | 145 | 4,261 | - |
| Longest contigs | 1,834,218 | - | - |
| No of scaffolds | - | 2,983 | 498 |
| Longest scaffold | - | 108,686 | - |
| Gaps within scaffolds | - | 459,015 | 236 |
| Number of pseudo-chromosomes | - | - | 14 |
| Amount of N's | 0 | 459,015 | 27,778 |
| Nb. of genes | 5,570 | 5,988 | 5,146 |
| Nb. of tRNAs | 40 | - | 46 |
| Nb. of putative transposable elements | 7 | - | 344 |
| Genome: BUSCO-apicomplexa_odb10 (n=446) | C:96.9 (S:94.6 - D:2.2) | C:93.3 (S:93.3 - D:0) | C:100 (S:100 - D:0) |
| Genome: BUSCO-plasmodium_odb10 (n=3642) | C:78.7 (S:75.4 - D:3.3) | C:66.4 (S:66.4 - D:0) | C:98.1 (S:98.0 - D:0.1) |
| Proteome: BUSCO-apicomplexa_odb10 (n=446) | C:90.8% (S:90.1% -D: 0.7%) | C:69.3% (S:68.8% -D:0.4%) | C:100 (S:100 - D:0) |
| Proteome: BUSCO-plasmodium_odb10 (n=3642) | C:68.0% (S:65.8% -D:2.2%) | C:41.4% (S:41.4% -D:0.0%) | C:98.8% (S:98.7% -D:0.1%) |

BUSCO scores (in %) for complete (C) orthologous gene sets, including those present in single-copy (S), and those duplicated (D).

Genome completeness was assessed using both the Apicomplexa (446 BUSCO groups) and Plasmodium (3,642 BUSCO groups) reference datasets, as no *Haemoproteus* species are currently represented in BUSCO. The Aplicomplexa dataset indicated high completeness (97% including 95% as single-copy orthologus), whereas lower scores (79%) were obtained with the *Plasmodium* dataset (**Table 1**). This may likely reflect evolutionary divergence and gene loss between the two genera.

The maximum likelihood phylogenetic tree of 12 species of the *Plasmodium* genus and 2 *Haemoproteus* species (**Figure 1**) estimated from 142 single-copy BUSCO orthologs (120,269 amino acids) was consistent with previous phylogenetic studies [2,5] The WW2 *H. majoris* linage branched as a sister species to *H. tartakovskyi* SISKIN1 lineage (with ∼25% divergence) in a clade outside the *Plasmodium* genus (**Figure 1**). Genetic divergence between the WW2 lineage and the 12 *Plasmodium* species ranged between 44% an 60%.

**Figure 1.**
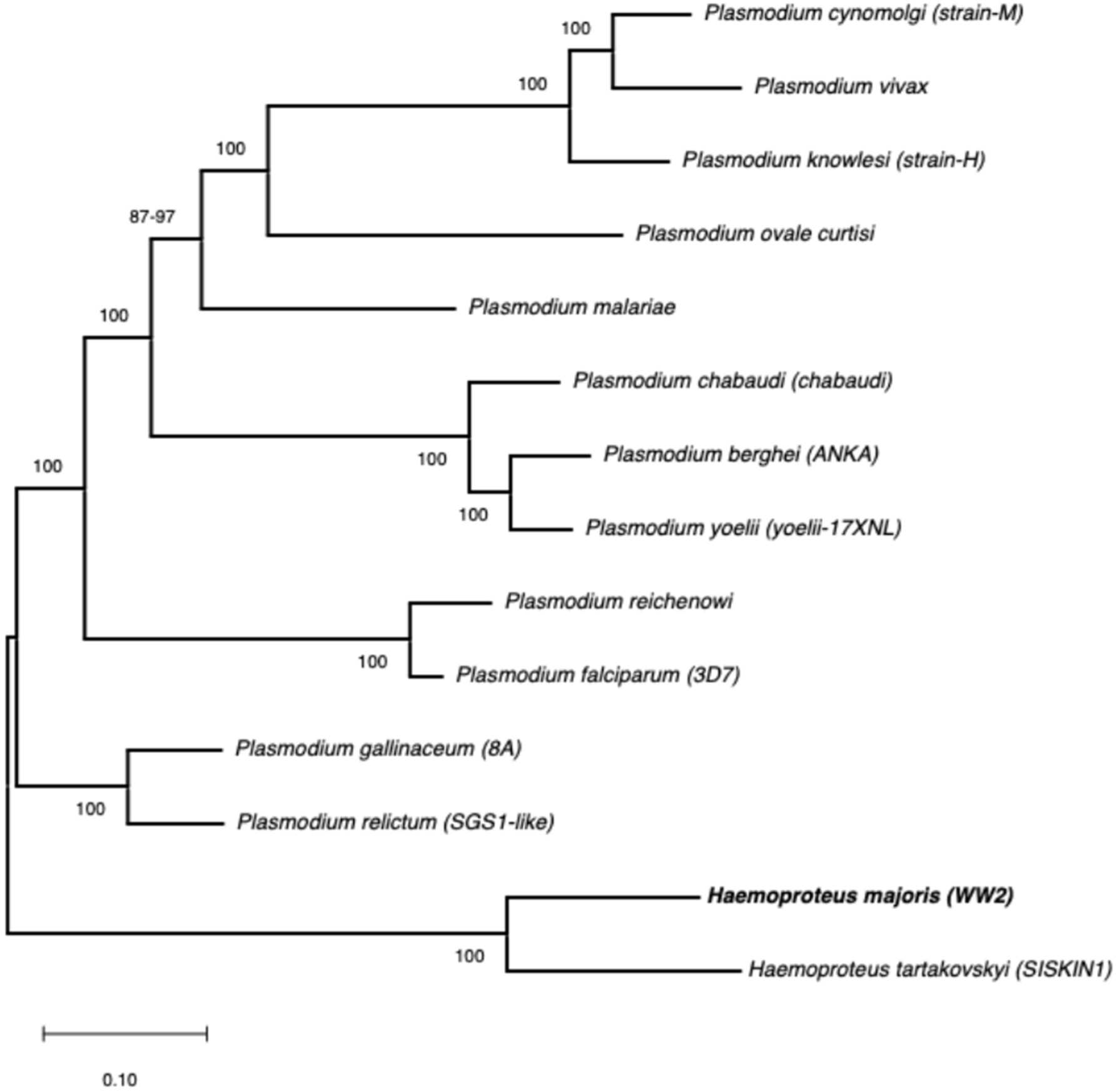
Phylogenetic relationships among major lineages of *Plasmodium* and *Heamoproteus* highlighing the newly generated *H. majoris* reference genome. The maximum likelihood (ML) phylogenetic tree, based on 142 single-copy BUSCO orthologs (*Apicomplexa database)* consisting in 120,269 amino acids, was estimated using the JTT+F+G+I model of amino acid evolution identified as the best-fitted model by MEGA12, based on the BIC values. The values at the node indicate node support (in percent) estimated with rapid adaptive bootstrap.

### Ortholog and lineage specific genes between *H. majoris* WW2 and *P. relictum* SGS1-like

During annotation, *Companion* pipeline [9] identified 296 putative pseudogenes, considerably more than the 40 *P. relictum* SGS1-like in PlasmoDB [6], all of which were retained for downstream analyses. Companion has clustered 5501 WW2 genes which are separated in to major groups, the shared orthologous groups with the reference and the orthologous groups within WW2. A total of 79.2% of WW2 orthologous protein-coding genes clustered into shared orthologous groups with SGS1-like, a higher proportion than the 60% reported for *H. tartakovskyi* SISKIN1 across a broader set of apicomplexan parasites [3]. Among shared orthologous clusters, 79.9% were present as single-copy genes in both genomes, and approximately 74.4% of the shared orthologous protein coding genes could be assigned a putative function (excluding « hypothetical protein » and « conserved protein »). The relatively high proportion of genes lacking functional annotation is consistent with patterns reported across haemosporidian genomes, where AT-rich composition reduces sequence similarity to other organisms.

The remaining 20.8% of the annotated protein-coding genes in WW2 were lineage specific, of which 79.7% occurred as single copy genes, consistent with observations in *H. tartakovskyi* [3]. Elucidating the origins and functions of these lineage-specific genes through ancestral state reconstruction would be a valuable avenue for future work.

### Ribosomal RNA (rRNA) and transfer RNA (tRNA) genes

Annotation of the *H. majoris* WW2 assembly identified one copy of the 5.8S gene and six copies of the 5S rRNA gene; the 18S and 28S rRNA genes were not detected. For comparison, the *H. tartakovskyi* SISKIN1 genome was reported to contain two copies of the 18S and two copies of 28S genes [3]. As Bohme *et al.* [2] did not annotate rRNA genes for either *P. relictum* SGS1-like or *P. gallinaceum* 8A [2], we queried PlasmoDB (Release 68, [6]) using BLAST and recovered between one and five copies of the 18S, 5.8S, 28S and 5S genes. The incomplete and inconsistent rRNA annotations across these genomes complicate direct comparisons, and may result from differences in assembly contiguity and annotation pipelines as much as genuine biological variation.

A reduced tRNA gene repertoire was obtained in our assembled genome of *H. majoris* WW2, with only 40 tRNA genes identified, consistent with the pattern observed across haemosporidians: 42 tRNA in SISKIN1 [6], and 46 in SGS1-like and 8A [2]. In the human malaria parasite *Plasmodium falciparum* 46 tRNA genes were identified, one per tRNA isoaceptor [7]. This reduced tRNA repertoire has led to the hypothesis that haemosporidians may rely on exogenous sources of tRNA. Supporting this, Bour *et al* [8] identified a surface protein in *P. falciparum,* the tRNA import protein (tRIP), capable of binding and importing host-derived tRNAs, representing the first known case of tRNA import in an eukaryote. Orthologs of *tRIP* are present in one copy in *P. relictum* SGS1-like, *P. gallinaceum* 8A, *H. tartakovskyi* SISKIN1 and in one pseudogenic copy in *H. majoris* WW2 [6].

The evolutionary implications of tRIP are particularly intriguing in the context of avian haemosporidians. In mammalian hosts, erythrocytes are anucleate and transcriptionally inert, so any imported tRNAs must originate from other nucleated host cells. Avian erythrocytes, however, are nucleated and retain an active transcriptional machinery raising the possibility that imported tRNAs may originate directly from the nucleated erythrocyte itself, a possibility that warrants future investigation.

### Multicopy gene families

Subtelomeric regions of Apicomplexan genomes are typically enriched by lineage-specific multicopy gene families [10]. In *Plasmodium*, these include well-characterised variant antigen families such as *var*, *rif*, and *stevor* in *P. falciparum* [7], *vir* and *PVpir* in *P. vivax* [11]. Previous avian *Plasmodium* sequences have identified *PIR-like*, *SURFIN*, *ETMB*, *RBP* and 6 *fam* genes two of which (*fam-g* and *fam-h*) are specific to avian parasites. Notably, none of these families were detected in *H. majoris* WW2 consistent with their absence in *H. tartakovskyi* SISKIN1. Both *Haemoproteus* genomes do, however, contain *ETRAMP* (early transcribed membrane protein) and *MSP* (merozoite surface protein) family members: one *ETRAMP* and seven *MSP* copies in WW2, and one *ETRAMP* and five *MSP* copies in SISKIN1 [6].

Orthology clustering identified 362 expanded gene families in WW2, the majority comprising fewer than 5 copies. The most striking expansion involves the 6-cysteine family, a protein involved in host cell invasion, immune evasion, which comprises 35 copies in WW2 versus 11 in SGS1-like. Several members show marked copy number variation: *P38* (17 copies in WW2 vs 1 in SGS1-like), *P92* (3 vs 1) and the related s48/47 domain-containing proteins (7 vs 0), while others are present as single conserved orthologs in both genomes (p230p, P12, P52, P36, P41, B9, P47 and one copy annotated “6-cysteine protein”).

### Transposable elements

Transposable elements (TEs) are notably absent from human and rodent *Plasmodium*, but have been reported in several avian haemosporidia species, including *P. relictum* SGS1-like, *P. gallinaceum* 8A and *H. tartakovskyi* SISKIN1 [12]. In *H. majoris* WW2, we identified 57 LTR retrotransposons (LTR-RTs) comprising one full-length LTR element, with 56 truncated elements and solo LTRs, all belonging to a single TE family, considerably fewer than the 344 LTR-RT copies detected in SGS1-like (Table 1). However, the apparent paucity of TEs in these genomes should be interpreted with caution, as their detection may be limited by the AT composition of these genomes, high sequence divergence from available TE libraries, and the absence of curated TE references for avian haemosporidians.

## CONCLUSION

In summary, we present the first high-quality assembled genome of *H. majoris* (lineage WW2), a prevalent and host generalist avian blood parasite. The resulting assembly represents a substantial improvement in contiguity over the sole previously published *Haemoproteous* genome, and constitutes a valuable addition to the scarce genomic resources available for avian haemosporidians. Comparative analyses reveal gene content and genomic architecture broadly consistent with other haemosporidian parasites, including a markedly reduced tRNA repertoire and the presence of a tRIP orthologue, a finding that raises intriguing questions about the sources of exogenous tRNA in parasites infecting nucleated avian erythrocytes. The detection of LTR retrotransposons further supports the emerging picture of avian haemosporidian genomes as distinctly different from their human and rodent *Plasmodium* counterparts. Given the basal phylogenetic position of avian haemosporidians, genomic resources from taxa such as *H. majoris* is essential for elucidating the evolutionary origins of key malaria parasite traits.

## MATERIALS AND METHODS

### Sample collection

The infected blood sample originated from a willow warbler (*Phylloscopus trochilus*) caught during autumn migration at Lake Krankesjön, Skåne, Sweden by one of the co-authors (OH). One aliquot of the blood was stored into SET-buffer for later DNA extraction and parasite identification. A second aliquot was snap frozen on dry ice and stored at -80°C prior to FACS-sorting.

### Parasite identification and isolation

The parasite was identified as *H. majoris* lineage WW2 using an established nested PCR-protocol resulting in the amplification of a 480 bp fragment of the *cytochrome b,* followed by Sanger sequencing [13]. This mitochondrial region is used as the gold standard for identification of avian haemosporidian parasites [14]. Prior to sequencing, parasite DNA was enriched by FACS-based sorting of parasite blood stages away from host cell nuclei. Briefly, parasite isolation was performed following Boissière *et al*. [15] from 40 µl blood. Propidium Iodide (PI)-stained samples were sorted on a FACSAria III (Becton Dickinson) cell sorter, and bird cell nuclei and parasite blood stages were discriminated based on their distinct scatter plots. Parasites were sorted into 96-well plates containing 3.5µl of PBS per well for subsequent whole-genome amplification.

### Whole genome amplification and sequencing

Whole genome amplification (WGA) was performed using the REPLIg Midi kit (Qiagen) following the manufacturer’s instructions. Both Nanopore and Illumina sequencing technologies were conducted by the MGX-Montpellier GenomiX platform. For Nanopore sequencing, 3144 ng of WGA-derived DNA was used to construct a library with the 1D Ligation Sequencing Kit (SQK-LSK110, Oxford Nanopore Technologies, ONT). Sequencing was performed on a MinION sequencer (ONT) with the MinKNOW software (v22.05.5), and quality control assessed out using PycoQC (v2.5.2) [16]. For Illumina sequencing, PCR-free libraries were prepared using the TruSeq DNA PCR-Free kit (Illumina, USA), and validated by Agilent Fragment Analyzer (High Sensitivity NGS kit) and qPCR (Roche Light Cycler 480). Pair-end sequencing (2 x 150bp) was performed on an Illumina NovaSeq 6000 (SP flow cell, 300 cycles). Demultiplexing and production of fastq files were performed with Illumina’s bcl2fastq software (v2.20.0.422) and sequencing quality was assessed with FastQC (v0.11.9) [17].

### Genome assembly

The assembly pipeline and parameters are available at https://github.com/mikafontaine/malaria_HaemoproteusWW2.git. Nanopore long-reads were first cleaned using NanoFilt (v2.8.0) [18], then assembled with FLYE (v2.8.3) [19] and the draft assembly was refined through one iteration of RACON (v1.4.2) [20] and one iteration of MEDAKA (v1.6.0) [21]. Further polishing was performed using the ILRA (Improvement of Long Read Assemblies pipeline) [22], which corrects and merges contigs, homopolymer tracks and circularizes plastids. Contaminated contigs were removed with Centrifuge/Recentrifuge (v1.0.4_beta/v1.3.2) [23,24]. Due to the size of the Illumina read files, short reads were downsampled using VariantBam (v1.4.4a) [25] retaining a maximum of 1000 reads per position prior to correction within ILRA. Homopolymer errors, single-base discrepancies and indels were corrected using 3 iterations of Pilon (v1.23) with the downsampled Illumina reads [26]. Assembly quality was assessed with BUSCO (v5.8.3) [27] and QUAST (v5.2.0) [28].

### Gene and transposable element annotations

Initial gene annotation of the unmasked WW2 genome was performed with *Companion* (v2.2.11, Release 63) [9], using *P. relictum* SGS1-like as reference. Orthology analysis was conducted with *OrthoFinder* (v2.5.4) [9,29], based on Diamond hits between translated protein sequences of the target and reference genomes. Protein completeness was assessed with BUSCO (v5.8.3) [27]. Comparative analyses with published *Plasmodium* and *Haemoproteous* genomes relied on *PlasmoDB* (Release 68).

As *Companion* identifies transposable elements (TEs) as genes, TEs were subsequently annotated using *EDTA* (v2.2.2) [30]. To date, only LTR gypsy-like retrotransposons have been identified in avian malaria (*Plasmodium* and *Heamoproteus*) [12]. Accordingly, repetitive sequences other than the LTR-RT retrotransposons (LTR-RTs) were excluded, as the remaining repeats likely reflect the AT-rich and repetitive nature of the genome. Positions of LTR-RTs identified by *Companion* and TE domains identified with *EDTA* overlapped, while LTR-RTs positions showed no overlap with annotated genes. LTR-RTs exceeding 500bp were masked using *BEDTools maskfasta* [31] (six TEs annotated as « gene » by Companion and not detected by EDTA have been used in the masking of the genome) and a second round of annotation was performed following the same procedure as above.

### Phylogenetic Analyses

Phylogenetic analyses of the newly generated WW2 *H. majoris* genome with other *Haemoproteus* and *Plasmodium* was based on single-copy BUSCO proteins from the apicomplexa_odb10 dataset (plasmodium_odb10.2024-01-08.tar.gz, https://busco.ezlab.org/).

BUSCO analysis was performed for WW2 and SISKIN1 and only BUSCO groups common to both assemblies were retained. From the plasmodium_odb10 dataset, 12 *Plasmodium* species were included (*P. berghei* ANKA, *P. gallinaceum*, *P. knowlesi* strain H, *P. reichenowi*, *P. vivax*, *P. malariae*, *P. chabaudi chabaudi*, *P. falciparum* 3D7, *P. relictum*, *P. yoelii yoelii* 17XNL, *P. ovale curtisi* and *P. cynomolgi* strain M), consistent with those used in previous avian malaria genome studies [2,5]. Out of the 446 BUSCO groups assessed, 142 on single-copy BUSCO orthologs were retained based on representation in at least 10 *Plasmodium* lineages as well as WW2 and SISKIN1.

Protein sequences from each individual BUSCO group were concatenated using Seqkit v2.10.1 [33], aligned with MAFFT v7.490 [34], and visualized with Geneious prime 2025.1.3. The maximum likelihood (ML) phylogenetic analysis was conducted using MEGA12 [35] on the complete protein sequence alignment of 120,269 amino acids. The best protein sequence substitution model (Jones-Taylor-Thornton – JTT+G+I+F) was identified using a ML procedure, based on the BIC and AICc scores. The ML phylogenetic tree was then estimated using this substitution model and the complete alignement. Fast adaptive bootstrap resampling was conducted to assess node supports.

## DATA AVAILABILITY

Raw sequence reads and the annotated genome were deposited in the European Nucleotide Archive (ENA, https://www.ebi.ac.uk/ena/browser/home) under BioProject PRJEB76951 and BioSample SAMEA115829188. Illumina paired-end and nanopore raw reads are available under accession numbers ERR14207237 and ERR13899816, respectively. The annotated genome is deposited under accession number ERZ27247697. The scripts and pipelines used in this project are available at https://github.com/mikafontaine/malaria_HaemoproteusWW2. The original assembly, the genome annotation, as well as an archive of the Github describing all the bioinformatic steps are also available via the IRD DataSud repository https://doi.org/10.23708/0QPMON.

## ACKNOWLEDGMENTS

SG acknowledges support from ANR EVOMALWILD (grant ANR- 17-CE35-0012).

